# MITO-Tag Mice enable rapid isolation and multimodal profiling of mitochondria from specific cell types *in vivo*

**DOI:** 10.1101/425454

**Authors:** Erol Can Bayraktar, Lou Baudrier, Ceren Özerdem, Caroline A. Lewis, Sze Ham Chan, Tenzin Kunchok, Monther Abu-Remaileh, Andrew L. Cangelosi, David M. Sabatini, Kivanc Birsoy, Walter W. Chen

**Affiliations:** Laboratory of Metabolic Regulation and Genetics, The Rockefeller University, New York City, NY 10065, USA; Whitehead Institute for Biomedical Research and Massachusetts Institute of Technology, Department of Biology, 9 Cambridge Center, Cambridge, MA 02142, USA; Howard Hughes Medical Institute, Department of Biology, Massachusetts Institute of Technology, Cambridge, MA 02139, USA; Koch Institute for Integrative Cancer Research, 77 Massachusetts Avenue, Cambridge, MA 02139, USA; Broad Institute of Harvard and Massachusetts Institute of Technology, 7 Cambridge Center, Cambridge, MA 02142, USA; Present address: Boston Combined Residency Program, Department of Pediatrics, Boston Children’s Hospital, 300 Longwood Avenue, Boston, MA 02115, USA

## Abstract

Mitochondria are metabolic organelles that are essential for mammalian life, but the dynamics of mitochondrial metabolism within mammalian tissues *in vivo* remains incompletely understood. While whole-tissue metabolite profiling has been useful for studying metabolism *in vivo*, such an approach lacks resolution at the cellular and subcellular level. *In vivo* methods for interrogating organellar metabolites in specific cell-types within mammalian tissues have been limited. To address this, we built on prior work in which we exploited a mitochondrially-localized 3XHA epitope-tag (“MITO-Tag”) for the fast isolation of mitochondria from cultured cells to now generate “MITO-Tag Mice.” Affording spatiotemporal control over MITO-Tag expression, these transgenic animals enable the rapid, cell-type-specific immunoisolation of mitochondria from tissues, which we verified using a combination of proteomic and metabolomic approaches. Using MITO-Tag Mice and targeted and untargeted metabolite profiling, we identified changes during fasted and refed conditions in a diverse array of mitochondrial metabolites in hepatocytes and found metabolites that behaved differently at the mitochondrial versus whole-tissue level. MITO-Tag Mice should have utility for studying mitochondrial physiology and our strategy should be generally applicable for studying other mammalian organelles in specific cell-types *in vivo*.

## INTRODUCTION

Mitochondria are membrane-bound organelles that house metabolic pathways that play critical roles in mammalian physiology. The mitochondrial matrix, the innermost mitochondrial compartment, is a unique metabolic space within the cell because of the presence of a distinct array of metabolic processes and the selective permeability of the inner mitochondrial membrane (1). Because of the unique metabolic landscape within the mitochondrial matrix, metabolomic approaches with organellar-level resolution can provide important insights into mitochondrial and cellular physiology that otherwise may be missed using whole-cell or whole-tissue analyses (2). Importantly, given that mitochondria have been implicated in a wide variety of pathologies, such as myocardial infarction and Parkinson’s Disease (1, 3), a complete understanding of how mitochondrial dysfunction contributes to these various disease states will likely require the study of mitochondrial metabolites *in vivo*.

We have previously developed a mitochondrial metabolomics workflow for cultured mammalian cells that utilizes a mitochondrially-localized 3XHA epitope-tag (“MITO-Tag”) to achieve the rapid isolation of mitochondria followed by metabolite profiling by liquid chromatography and mass spectrometry (LC/MS) (2, 4). Utilizing this methodology, we were able to observe changes in matrix metabolism that were not appreciated using whole-cell metabolomics. Similar approaches from our laboratory for studying lysosomes in cultured cells have also revealed the dynamics of lysosomal metabolism during different cellular states (5, 6). Because mammalian physiology can often be better modeled *in vivo* rather than in cultured cells, we wished to implement our workflow for studying mitochondria in mice. Of particular importance, mitochondrially-targeted epitope-tags would allow for the rapid isolation of mitochondria from specific cell-types in complex tissues without the need for cell-sorting, thus improving the speed of the workflow and reducing distortion of the mitochondrial metabolite profile (7). This would be useful for cell-types such as neurons, as the state of these cells can be particularly vulnerable to distortions during traditional cell-sorting methods due to their frailty and sprawling cellular architecture (8). To date however, no such methodology exists for rapidly isolating mitochondria with cell-type-specificity from mammalian tissues *in vivo* (9-14).

To that end, we generated “MITO-Tag Mice,” *Rosa26* knock-in mice that express the MITO-Tag only in cells that express Cre recombinase, as a result of a loxP-STOP-loxP (LSL) cassette. This strategy allows for rapid isolation of mitochondria with cell-type-specificity from mammalian tissues as long as one has an appropriate promoter driving expression of Cre recombinase (15). Note that the MITO-Tag is the 3XHA-EGFPOMP25 or HA-MITO construct described previously (2, 4). We show that these MITO-Tag Mice allowed for the rapid immunopurification and both proteomic and metabolomic analyses of mitochondria from hepatocytes *in vivo*. Using both targeted and untargeted metabolomics, we uncovered changes in a diverse array of mitochondrial metabolites during fasted and refed conditions, thus shedding light on how hepatocyte mitochondria behave during these two important nutritional states. We also identified metabolites that behave differently when assessed at the mitochondrial versus whole-tissue level, thus underscoring the importance of mitochondrial metabolomics. Taken together, our work demonstrates the utility of MITO-Tag Mice for studying mitochondrial biology *in vivo* and offers a framework for studying other mammalian organelles in specific cell-types within complex tissues.

## RESULTS

To enable rapid isolation of mitochondria from specific cell-types *in vivo*, we generated “MITO-Tag Mice” by knocking our HA-MITO construct and an upstream LSL cassette into the *Rosa26* locus (Fig. 1*A*). In the absence of Cre recombinase, the LSL cassette will prevent the production of the HA-MITO protein and anti-HA immunopurifications will not yield mitochondria. However, once present, Cre recombinase will excise the LSL cassette and thus enable epitope-tagging and rapid isolation of mitochondria. Importantly, by selecting the appropriate promoters for controlling the expression of Cre recombinase, one can restrict epitope-tagging of mitochondria to specific cell-types and consequently immunoisolate mitochondria with cell-type-specificity from a piece of tissue without the need for cell sorting.

**Fig. 1.**
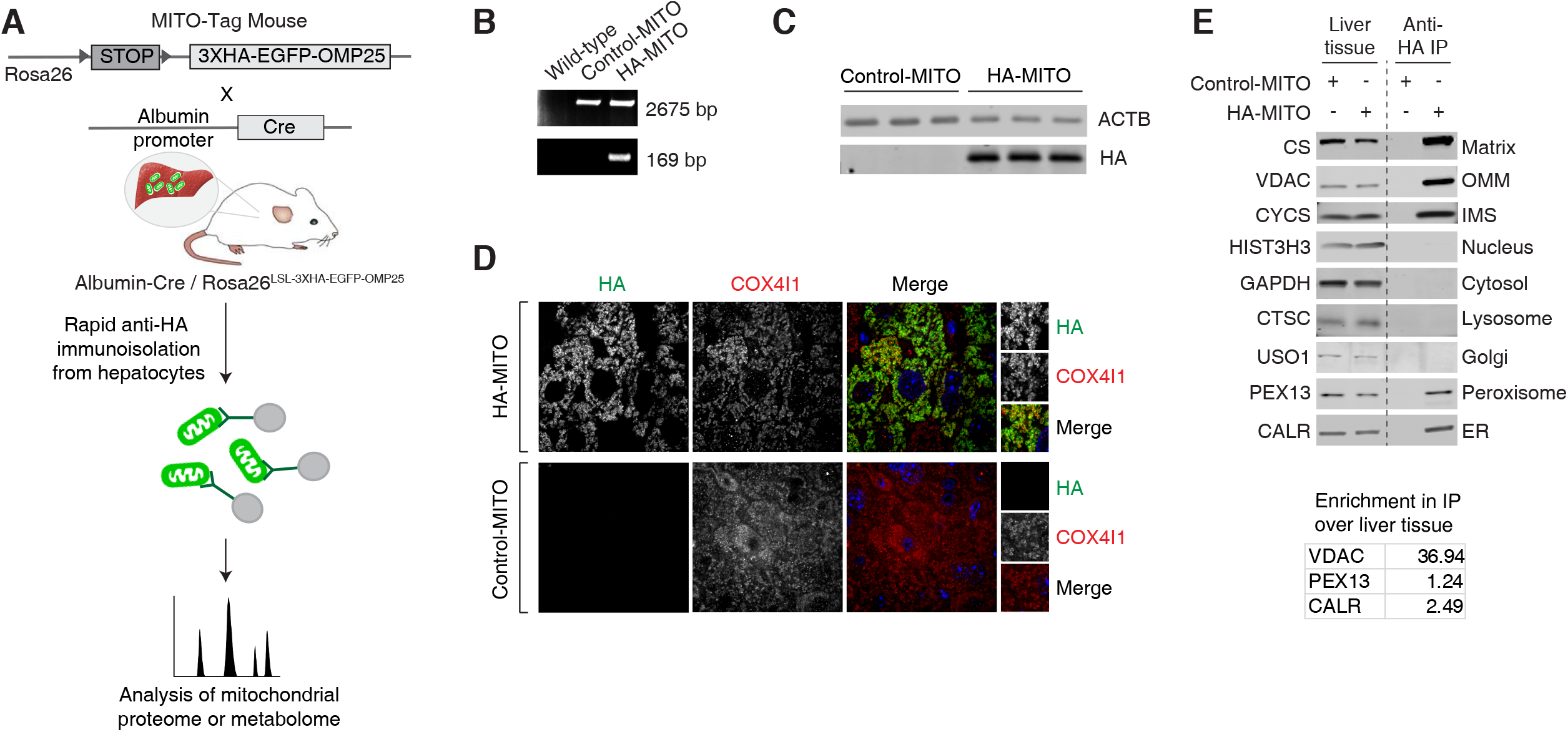
Characterization of MITO-Tag Mice and associated *in vivo* workflow. (*A*) Schematic demonstrating the design of the MITO-Tag Mice and how they can be utilized for rapid, cell-type-specific isolation and multimodal analysis of mitochondria *in vivo*. MITO-Tag Mice contain the *3XHA-EGFP-OMP25* (*HA-MITO*) gene knocked into the *Rosa26* locus. The expression of this gene is dependent on the presence of Cre recombinase due to an upstream loxP-STOP-loxP (LSL) cassette. Mating MITO-Tag Mice with mice expressing Cre under the control of certain promoters, such as the Albumin promoter as in this study, allows for cell-type-specific expression of the *HAMITO* gene and, consequently, allows for the rapid, cell-type-specific isolation of mitochondria *in vivo* (mitochondrial isolation takes ~10 min once liver tissue has been homogenized). Because “KPBS,” the mitochondrial isolation buffer utilized in our workflow, is compatible with mass spectrometric analyses, the isolated mitochondria can subsequently be interrogated with either proteomics or metabolomics. The actual immunopurification workflow used here is overall similar to that used in our prior work (2, 4), however, it is important to note that the material inputs in this work are liver tissue from *Rosa26^LSL-3XHA-EGFP-OMP25/+^* mice (Control-MITO mice) and *Alb-Cre^+/-^*, *Rosa26^LSL-3XHA^EGFP-OMP25/+* mice (HA-MITO mice) and no measurements of mitochondrial volume were done in this study because we performed relative rather than absolute quantification of metabolites. Arrowheads indicate loxP sites. (*B*) Genomic PCR analysis of liver tissue taken from mice with the indicated genotypes. The upper (2675 bp) and lower (169 bp) bands indicate the presence of the desired knock-in at the *Rosa26* locus and successful Cre-mediated excision of the floxed region, respectively. (*C*) Immunoblot analysis of liver tissue taken from mice with the indicated genotypes. ACTB was used as a loading control. (*D*) Immunofluorescence of liver tissue taken from mice with the indicated genotypes. HA (green), COX4l1 (red), and Hoescht (blue) signals are shown. (*E*) Immunoblot analysis of liver tissue and the anti-HA immunoprecipitates (anti-HA IP) from mice with the indicated genotypes. The line indicates that two parts of the same membrane have been brought together. The names of the protein markers used and the corresponding subcellular compartments appear to the left and right of the immunoblots, respectively. Matrix, mitochondrial matrix; OMM, outer mitochondrial membrane; IMS, mitochondrial intermembrane space; Golgi, Golgi complex; ER, endoplasmic reticulum. The enrichment in the anti-HA IP, relative to liver tissue, of the markers VDAC, PEX13, and CALR was quantified and is shown in the inset table. All mice used for these experiments were fed *ad libitum*. For all figure panels, the following abbreviations are used: Control-MITO, Control-MITO mice; HA-MITO, HA-MITO mice.

To begin our studies utilizing the MITO-Tag Mice, we chose to examine mitochondria within hepatocytes, as these cells respond to a range of interesting physiological stimuli and possess mitochondria that house a diverse array of metabolic reactions. To specifically isolate mitochondria from hepatocytes, we mated MITO-Tag Mice to transgenic animals expressing Cre recombinase under the control of the Albumin promoter (Fig. 1*A*). Subsequent progeny could then be used for the rapid immunopurification of mitochondria from hepatocytes, with mitochondrial isolation taking ~ 10 min once the liver tissue had been homogenized. Because the mitochondrial isolation buffer used in our workflow (i.e., “KPBS”) is compatible with mass spectrometric analyses, the isolated mitochondria could then be characterized using either proteomic or metabolomic analyses. The immunopurification workflow was generally similar to that used in our prior studies with two primary differences (2, 4): firstly, the material inputs in this work were liver tissue from *Rosa26^LSL-3XHA-EGFP-OMP25/+^* mice (Control-MITO mice) and *Alb-Cre^+/-^*, *Rosa26^LSL-3XHA-EGFP-OMP25/+^* mice (HA-MITO mice); and secondly, no measurement of matrix volume was done because we performed relative rather than absolute quantification of metabolites.

Importantly, MITO-Tag Mice behaved as expected when assessed using several orthogonal methods. Utilizing genomic PCR and immunoblot analyses of liver tissue, we observed that excision of the STOP codon and production of the HA-MITO protein only occurred in the presence of Cre recombinase, respectively (Fig. 1 *B* and *C*). Immunofluorescent examination also revealed that the HA-MITO protein co-localized with COX4l1, a mitochondrial marker (Fig. 1*D*). Cells lacking HA-MITO protein in the images could either not be hepatocytes and thus lack Albumin promoter activity or be a hepatocyte that did not express the HA-MITO gene, which we observed occasionally. Immunoblot analysis of the HA-MITO immunoprecipitate (HA-MITO IP) revealed not only substantial enrichment of mitochondria relative to multiple non-mitochondrial, subcellular compartments, but also minimal organellar contamination of the Control-MITO immunoprecipitate (Control-MITO IP) (Fig. 1*E*). Consistent with the fact that mitochondria can form contacts with the endoplasmic reticulum and peroxisomes in certain cell types (16, 17), it is not surprising that our rapid, gentle workflow did not remove all traces of these two organelles, although mitochondrial markers were still highly enriched in the immunoprecipitate (IP) by comparison (see table in Fig. 1*E*).

We next performed proteomic and metabolomic analyses on the IP material from our *in vivo* workflow. From the 1204 proteins in our primary proteomics data, we identified 511 proteins that met a stringent enrichment criterion of having mean protein signal in the HA-MITO IPs (i.e., isolated mitochondria) that was at least five times greater than that in the Control-MITO IPs (i.e., background); of these 511 proteins with signal considerably above background, 75.9% of them were found in MitoCarta2.0 (Fig. 2*A* and Table S1) (18). Our proteomic analysis also provided evidence supporting the cell-type-specificity of our *in vivo* workflow (Table S1). We were able to detect BDH1, an enzyme required for synthesis of the ketone body 3-hydroxybutyrate and known to be present within hepatocyte mitochondria (19), at ~15-fold signal above background. In addition, BCL2, a mitochondrial protein present in lymphocytes but much less so in hepatocytes (20, 21), was absent from our primary proteomics data. We chose to look for a mitochondrial protein in lymphocytes because circulating lymphocytes can in principle be present within the liver tissue as not all blood is washed out of the tissue prior to homogenization, but we would not expect to isolate lymphocyte mitochondria since lymphocytes should not contain Cre recombinase when its expression is dictated by the Albumin promoter. We also carried out metabolomic analyses of the IP material from our *in vivo* workflow. Using lipidomics, we were able to detect numerous species of cardiolipin, a well-known class of mitochondrial lipids, in the HA-MITO IPs at levels that were greater than background and met our criteria for being mitochondrial (Fig. 2*B* and Table S2). Likewise, using polar metabolomics, we found that the HA-MITO IPs had known mitochondrial metabolites, such as NAD, NADH, NADP, NADPH, coenzyme A, alpha-ketoglutarate, ATP, 3-hydroxybutyrate, and saccharopine, at levels above background and meeting our criteria for being mitochondrial (Fig. 2*C* and Table S3). We also found that metabolites not expected to be in mitochondria, such as cystine (lysosome) and sedoheptulose 7-phosphate (cytosol), were undetectable in both Control-MITO and HA-MITO IPs. Collectively, these data demonstrate that our *in vivo* workflow successfully isolates mitochondria and can be utilized for proteomic, lipidomic, and polar metabolomic analyses.

**Fig. 2.**
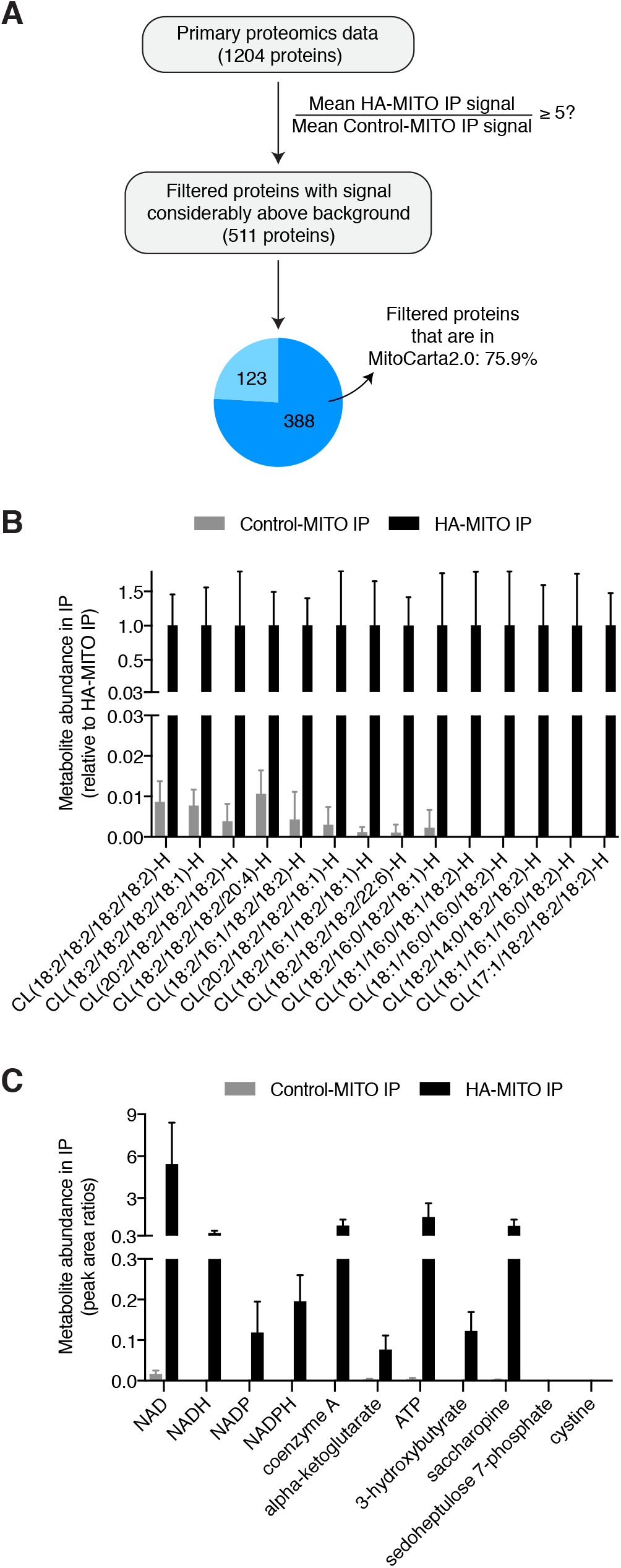
Mitochondria isolated with MITO-Tag Mice can be used for multiple downstream analyses and possess both protein and metabolite features characteristic of mitochondria. (*A*) Flowchart of how proteomics analysis was done. In brief, the 1204 proteins in our primary proteomics data were filtered based on a stringent enrichment criterion of having a mean pseudonumbered protein signal in the HA-MITO IPs that was at least five times greater than that in the Control-MITO IPs. Of these 511 proteins with signal considerably above that of the Control-MITO IPs (i.e., the background), 75.9% of them were found in MitoCarta2.0 by crossreferencing the NCBI gene IDs, with the respective numerical breakdown indicated in the pie chart wedges
(18). Mice used for these experiments were fed *ad libitum*. See also Table S1. (*B*) Lipidomic analysis of IPs for cardiolipin species. Negative lipid ion peak area signals for various cardiolipins (CLs) are used to generate this graph. Control-MITO IPs (n = 4 mice), HA-MITO IPs (n = 4 mice). Mice were fasted for 12 h and then refed for 4 h. Data are shown as means with SDs. All CLs presented here were categorized as grade A and satisfy the criteria of being considered mitochondrial, as defined in the Materials and Methods. See also Table S2. (*C*) Targeted metabolomic analysis of IPs for various polar metabolites. Shown are metabolite peak area ratios, which are calculated by dividing the metabolite peak areas by internal standard peak areas, for the Control-MITO and HAMITO IPs. NAD, NADH, NADP, NADPH, coenzyme A, alpha-ketoglutarate, ATP, 3-hydroxybutyrate, saccharopine are known mitochondrial metabolites, whereas cystine (lysosome) and sedoheptulose 7-phosphate (cytosol) are markers for extra-mitochondrial compartments. Control-MITO IPs (n = 4 mice), HA-MITO IPs (n = 4 mice). Mice were fasted for 12 h and then refed for 4 h. Data are shown as means with SDs. With the exception of cystine and sedoheptulose 7-phosphate, all metabolites shown here meet the criteria of being considered mitochondrial, as defined in the Materials and Methods. See also Table S3.

Because the polar metabolome of mitochondria is generally more prone to distortion during mitochondrial isolation than the proteome and lipidome are, we next leveraged the rapidity of our cell-type-specific workflow to study how polar mitochondrial metabolites behave in hepatocytes during fasted and refed conditions, two important nutritional states. Control-MITO and HA-MITO mice were fasted for 12 h overnight and then either fasted or refed for 4 h, after which we interrogated polar metabolites from hepatocyte mitochondria and whole liver tissue (Fig. 3*A*). To verify that our fasting and refeeding conditions were having the intended effects, we confirmed that there was greater phosphorylation of ribosomal protein S6 in the livers of refed Control-MITO and HA-MITO mice (Fig. 3*B*). Immunoblot analyses revealed that the immunopurifications generally behaved similarly during fasting and refeeding conditions (Fig. S1), with similar patterns of mitochondrial enrichment as immunopurifications performed with mice that were fed *ad libitum* (Fig. 1*E*). Targeted, polar metabolomic analyses of liver tissue and hepatocyte mitochondria revealed changes across a diverse range of metabolites (Fig. 3*C* and Table S3). A reassuring change we observed was increased acetyl-CoA levels in hepatocyte mitochondria isolated from animals that were fasted but not refed, which is likely a result of increased fatty acid oxidation within the mitochondrial matrix during fasted conditions (Fig. 3*D* and Table S3). Mitochondrial coenzyme A levels were elevated in fasting conditions as well, potentially reflecting increased rates of ketogenesis within mitochondria. We also observed that mitochondrial methylcitrate was considerably elevated when mice were fasted but not refed, which may be secondary to increased amounts of mitochondrial propionyl CoA. In the fasted state, there were also considerably elevated levels of mitochondrial GMP, which may indicate altered energy charge dynamics within the matrix, and increased amounts of mitochondrial glutamine, which may reflect decreased matrix glutaminolysis. Taken together, our data demonstrate the utility of our *in vivo* workflow for interrogating changes in polar mitochondrial metabolites during different physiological states.

**Fig. 3.**
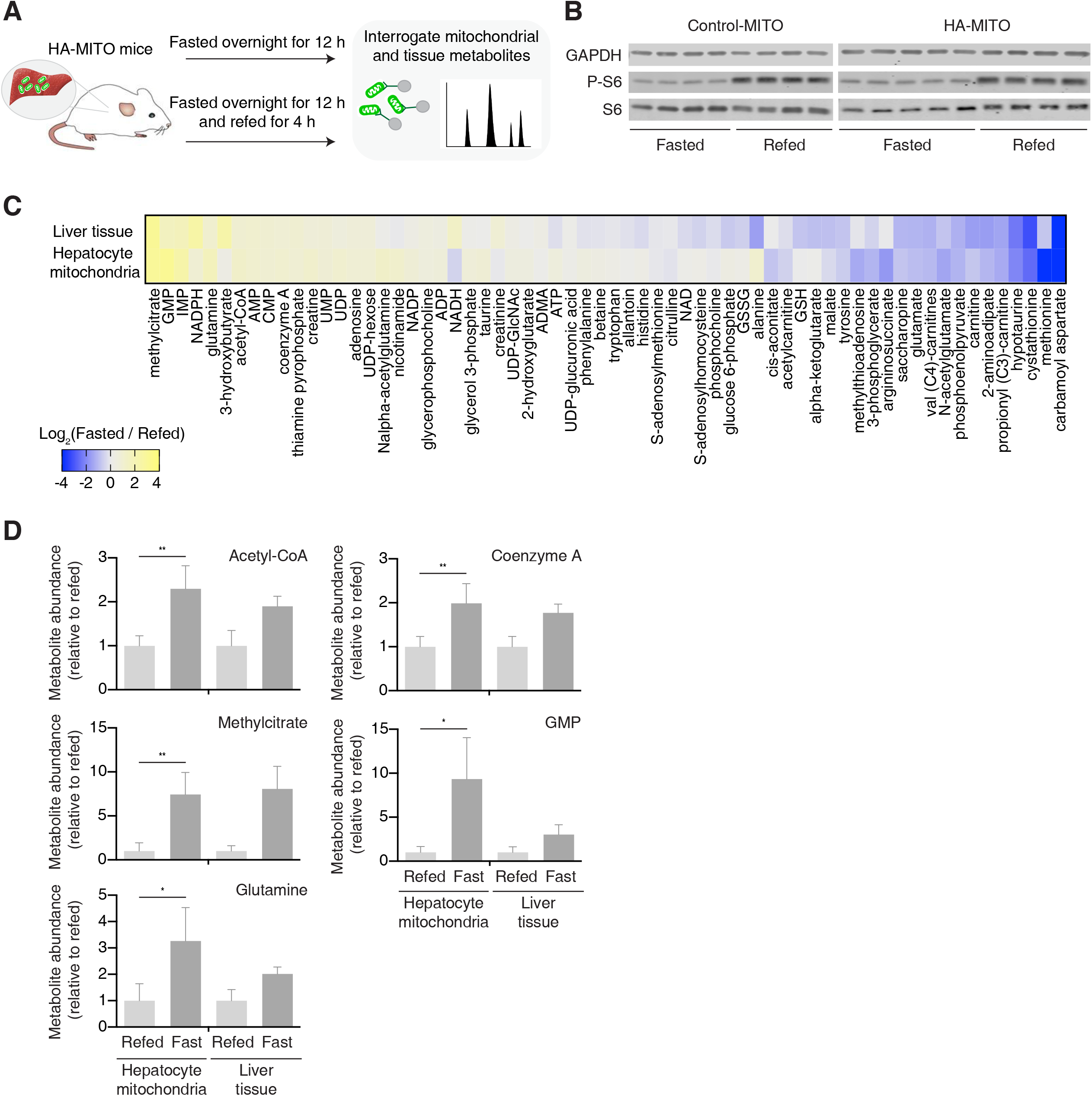
*In vivo* dynamics of hepatocyte mitochondrial metabolites during fasting and refeeding. (*A*) Schematic demonstrating general design of fasting and refeeding experiment. In brief, Control-MITO and HA-MITO mice were fasted overnight for 12 h and then either fasted or refed for another 4 h before liver tissue was harvested for mitochondrial or whole-tissue polar metabolomics. (*B*) Immunoblot analysis of liver tissue taken from Control-MITO and HA-MITO mice during fasted and refed conditions. Protein names appear to the left of the membrane. P-S6, phospho-S6 [Ser240/244]. GAPDH was used as a loading control. (*C*) Heat map of changes in metabolites as assessed at the mitochondrial and whole-tissue level. The data are presented as the log_2_-transformed mean fold change (fasted / refed). For a metabolite to be included in this heat map, it had to be considered mitochondrial in at least the fasted or refed state. See Materials and Methods for criteria for being considered mitochondrial. See also Table S3 for more detail. (*D*) Comparison of select metabolites during fasted and refed conditions. Metabolite peak area ratios, which are calculated by dividing the metabolite peak areas by internal standard peak areas, are used to generate these graphs. See also Table S3. Data are presented as means with SDs. *p < 0.05, **p < 0.01 as determined by unpaired, parametric, two-tailed student’s *t* test. For all panels, fasted Control-MITO mice (n = 4), fasted HA-MITO mice (n = 5), refed Control-MITO mice (n = 4), refed HA-MITO mice (n = 4).

To complement our targeted studies on polar mitochondrial metabolites, we also incorporated untargeted metabolomics into our *in vivo* workflow. From these untargeted analyses, we found several hits that had indeterminate chemical identities and significantly different abundances within the HA-MITO IPs during fasted and refed conditions (Fig. 4*A* and Table S4). Using authentic chemical standards that were commercially available, we successfully identified several of these hits to be Oadipoylcarnitine, O-succinylcarnitine, and 4-guanidinobutyric acid. Although the differences in O-adipoylcarnitine levels within mitochondria during fasted and refed conditions only trended towards statistical significance when peak areas were manually integrated and appropriately adjusted, the untargeted analysis was still fruitful as it allowed us to determine that, under both nutritional states, O-adipoylcarnitine was present in the HA-MITO IPs above background and at levels meeting our criteria for being considered mitochondrial (Fig. 4*B* and Table S4). We did find however that the differences in mitochondrial O-succinylcarnitine and 4-guanidinobutyric acid during fasted and refed conditions remained statistically significant after manual integration and appropriate adjustment of peak areas (Fig. 4*C* and Table S4), and it will be interesting to see future work decipher the mechanisms underlying these metabolic changes. Taken together, these data demonstrate how untargeted metabolomics can be used with our *in vivo* workflow to study mitochondrial metabolites without knowledge of their molecular identities beforehand.

**Fig. 4.**
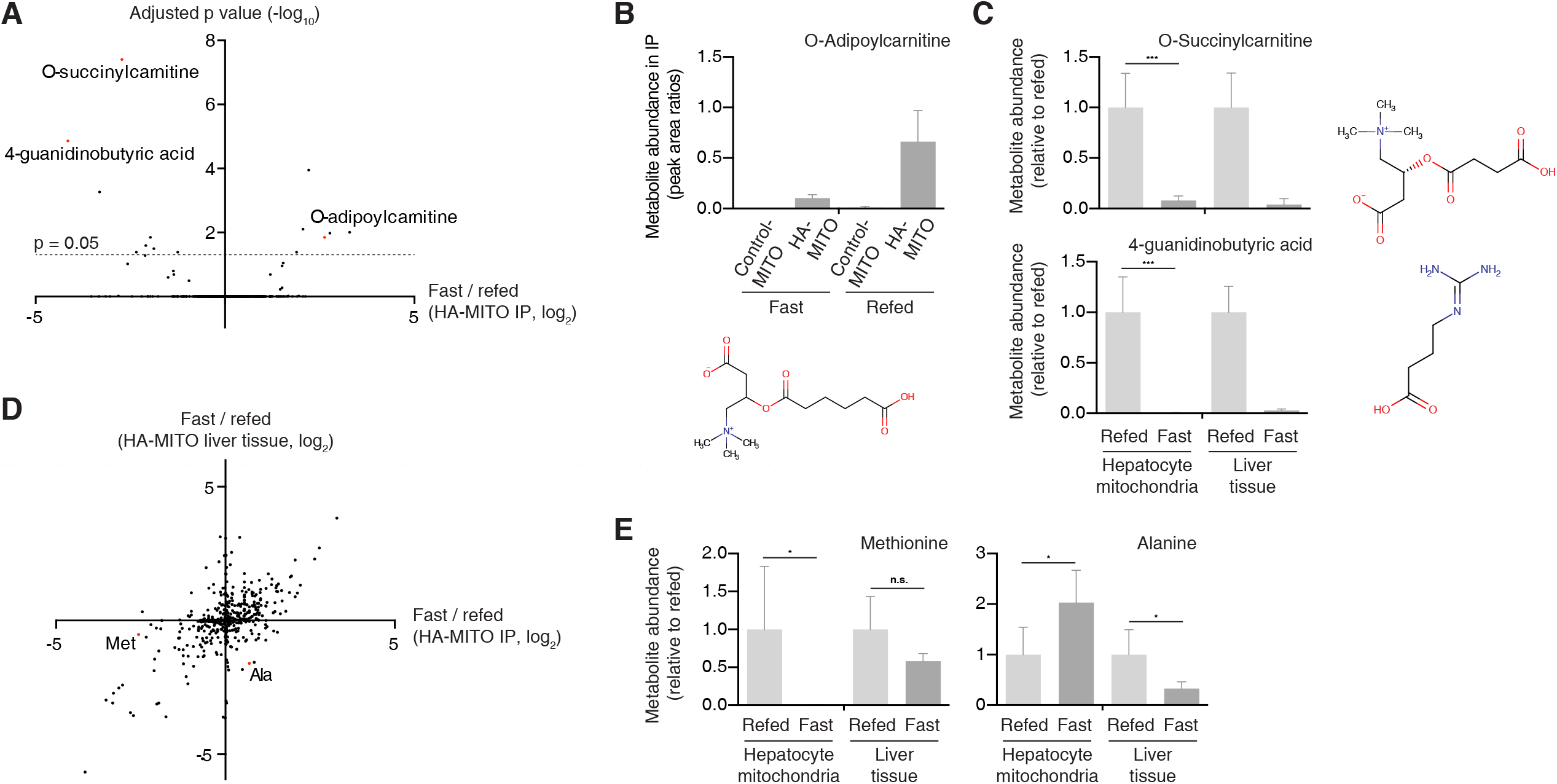
Interrogation of mitochondrial metabolites during fasting and refeeding using untargeted metabolomics. (*A*) Scatter plot of untargeted features in the HAMITO IPs (i.e., isolated mitochondria). The data presented here are the log_2_-transformed fold changes (median fasted HA-MITO IP signal / median refed HA-MITO IP signal). Ad hoc adjusted p values (Benjamini-Hochberg method) are presented here and the line indicates where the adjusted p value = 0.05. All points above the line are considered significant. Features that were subsequently validated using authentic chemical standards are highlighted in red and indicated by their metabolite name. See also Table S4 for more detail. (*B*) Metabolomic analysis of IPs for O-adipoylcarnitine. Shown are metabolite peak area ratios, which are calculated by dividing the metabolite peak areas by internal standard peak areas, for the Control-MITO and HA-MITO IPs during fasted and refed conditions. O-adipoylcarnitine meets the criteria of being mitochondrial in both fasted and refed conditions, as described in the Materials and Methods. The chemical structure of O-adipoylcarnitine is shown below the bar graph. Data are shown as means with SDs. See also Table S4. (*C*) Metabolomic interrogation of O-succinylcarnitine and 4-guanidinobutyric acid during fasted and refed conditions. Metabolite peak area ratios, which are calculated by dividing the metabolite peak areas by internal standard peak areas, are used to generate these graphs. See also Table S4. Data are shown as means with SDs. ***p < 0.001 as determined by unpaired, parametric, two-tailed student’s *t* test. The chemical structures of the respective metabolites are shown to the right of each bar graph. (*D*) Scatter plot of untargeted features when assessed on the mitochondrial versus whole-tissue level. The data presented here are the log_2_-transformed fold changes for the HA-MITO IP samples (median fasted HA-MITO IP signal / median refed HA-MITO IP signal) and the HA-MITO whole-tissue samples (median fasted HA-MITO whole-tissue signal / median refed HA-MITO whole-tissue signal). Methionine and alanine are highlighted in red as the behavior of these two metabolites was confirmed during our targeted analysis. See Tables S3 and S4 for more detail. (*E*) Metabolomic interrogation of methionine and alanine during fasted and refed conditions. Metabolite peak area ratios, which are calculated by dividing the metabolite peak areas by internal standard peak areas, are used to generate these graphs. See also Table S3. Data are shown as means with SDs. *p < 0.05 as determined by unpaired, parametric, two-tailed student’s *t* test. n.s., not statistically significant. For all panels, fasted Control-MITO mice (n = 4), fasted HA-MITO mice (n = 5), refed Control-MITO mice (n = 4), refed HA-MITO mice (n = 4).

Perhaps the most interesting metabolites identified by direct profiling of organelles are those that have significant discordance in their behaviors when assessed using organellar versus whole-tissue metabolomics. To that end, we also examined which untargeted metabolomic features behaved differently in the HA-MITO IPs versus in whole-tissue during fasted and refed conditions (Fig. 4*D* and Table S4). Our untargeted analysis identified methionine and alanine as two metabolites that exhibited the desired behavior. This was reassuring as the changes seen for these two metabolites were concurrently validated during our targeted analysis (Fig. 4*E* and Table S4). Interestingly, there were considerably reduced levels of methionine within mitochondria when mice were fasted but not refed, but methionine levels were not significantly different at the whole-tissue level. This may reflect that mitochondrial pools of methionine are less well-buffered during fasting than those in other compartments that contribute to the whole-tissue signal, such as the cytosol. With regards to alanine, we observed that fasted mice had elevated alanine within mitochondria but decreased levels of the amino acid at the whole-tissue level. The increase in mitochondrial alanine could be a result of decreased pyruvate dehydrogenase activity, increased mitochondrial pyruvate, and subsequent equilibration between pyruvate and alanine pools during fasting. The decreased amounts of whole-tissue alanine could be a reflection of an increased gluconeogenic drive to convert alanine to pyruvate within the cytosol. Collectively, these data demonstrate the value of our *in vivo* workflow, when used in conjunction with untargeted metabolomics, for taking an unbiased approach to studying mitochondrial metabolites during different physiological states.

## DISCUSSION

Methods for interrogating organellar metabolites with cell-type-specificity *in vivo* have thus far been limited in mammalian systems (9-14). Building on our prior methodology using epitope-tags for fast isolations of mitochondria from cultured cells (2, 4), we have generated MITO-Tag Mice that allow for the rapid, cell-type-specific immunopurification and multimodal analysis of mammalian mitochondria *in vivo*. By combining our *in vivo* workflow for rapid isolation of mitochondria with metabolomics, we uncovered changes in a diverse array of hepatocyte mitochondrial metabolites during fasted and refed conditions via targeted and untargeted approaches. Of note, we were able to identify metabolites, such as methionine and alanine, that behaved differently during fasted and refed conditions when assessed using mitochondrial versus whole-tissue metabolomics, thus underscoring the importance of examining metabolites at the organellar level.

We believe that, in addition to their utility for assessing mitochondrial metabolism *in vivo*, these MITO-Tag Mice will be a widely useful tool for studying mitochondrial physiology in mammalian systems. Multimodal analysis of mitochondria from specific cell types will be important for trying to dissect mitochondrial heterogeneity in complex tissues, such as the brain and kidney. Furthermore, the ability to rapidly isolate mitochondria from specific cell-types without the need for cell sorting and long centrifugation steps can reduce the magnitude of distortion of the original mitochondrial state, as well as significantly decrease the time and effort needed to acquire the desired mitochondrial material, which can be useful for not just metabolomic but also proteomic work by better preserving the native proteomic mitochondrial profile and the interactions of proteins that peripherally associate with mitochondrial membranes (22). Of note, the design of our *in vivo* workflow is such that it can also be combined with other downstream studies aside from those utilized here, such as genomic and transcriptomic analyses. We thus believe that these MITO-Tag Mice will have utility in the study of mitochondria, and feel that this work can serve as a general strategy for the cell-type-specific interrogation of other mammalian organelles *in vivo*.

## MATERIALS AND METHODS

### Reagents

Pierce Anti-HA magnetic beads (Cat: 88837)

Lucigen QuickExtract DNA Extraction Solution (Cat: QE09050)

Miltex Biopsy Punch with Plunger, ID 4.0 mm, OD 4.36mm (Ted Pella Cat: 15110-40)

Self Healing Cutting Mat, 3.5” X 5.5” (Ted Pella Cat: 15085)

VWR PowerMax Dual Shaft Laboratory Mixer (Cat: 14215-266)

Pierce BCA Protein Assay (Cat: 23224)

Roche cOmplete Mini EDTA-free protease inhibitor (Cat: 11836170001)

Roche PhosSTOP phosphatase inhibitor (Cat: 4906845001)

Omni International Bead Ruptor 24 Bead Mill Homogenizer (Cat: 10032-376)

Omni BR-Cryo Cooling Unit (Cat: 19-8005)

Cambridge Isotope Laboratories Metabolomics Amino Acid Mix Standard (Cat: MSK-A2-1.2)

Thermo Fisher Scientific Hoescht 33342 (Cat: H3570)

### Antibodies

For immunoblotting, the antibodies used were as follows: antibodies against CS (14309), VDAC (4661), HIST3H3 (4499), CALR (12238), S6 (2217) and P-S6 [Ser240/244] (2215) were purchased from Cell Signaling Technology; the antibody against CYCS was purchased from Thermo Fisher Scientific (MA511674); the antibody against GAPDH was purchased from GeneTex (GTX627408); the antibody against CTSC was purchased from Santa Cruz Biotechnology (sc-74590); the antibody against USO1 was purchased from BD Biosciences (612260); the antibody against PEX13 was purchased from MilliporeSigma (ABC 143). For immunofluorescence, the antibodies used were as follows: antibody against the HA epitope (3724) was purchased from Cell Signaling Technology; the antibody against COX4I1 was purchased from Abcam (ab14744). The Alexa Fluor 488-conjugated donkey anti-rabbit IgG (A-21206) and the Alexa Fluor 568-conjugated donkey anti-mouse IgG (A10037) were purchased from Thermo Fisher Scientific.

### MITO-Tag Mice

MITO-Tag Mice (*Rosa26^LSL-3XHA-EGFP-OMP25^* mice) were generated by first constructing a gene targeting plasmid using a CAG-driven R26TV LSL backbone that was gifted by Thales Papagiannakopoulos (Addgene: 64812). Gene blocks with the *HA-MITO* gene and homologous arms of 1 kb to the target site in the *Rosa26* locus were cloned into the vector with Gibson assembly. LoxP sites flanking the STOP codon were designated upstream of the *HA-MITO* gene. The construct was electroporated into Cy2.4 ES cells (B6(Cg)-Tyr<c2J> genetic background). Positive clones were identified by Southern blot analysis of the neomycin cassette. ES cells were injected into blastocysts and chimeras were screened based on white/black coat color selection and the strain was maintained heterozygous.

For the purposes of this study, we designated *Rosa26^LSL-3XHA-EGFP-OMP25/+^* mice as Control-MITO mice and *Alb-Cre^+/-^*, *Rosa26^LSL-3XHA-EGFP-OMP25/+^* mice as HA-MITO mice.

We generated HA-MITO mice by mating MITO-Tag Mice with mice expressing Cre recombinase under the control of the Albumin promoter (B6.Cg-Speer6-ps1Tg(AlbCre)21Mgn/J mice, purchased from The Jackson Laboratory, stock number: 003574) (23). PCR genotyping for alleles containing the *HA-MITO* and *Alb-Cre* genes were performed on isolated genomic DNA from mouse tails with following primers.

MITO-Tag_F: GGTCACCACACTCACCTATGGCGTACAGT

MITO-Tag_R: CCTTGCTCAGTGCAGATTGCGTGCTCAG

Alb-Cre_F: CTGCCACGACCAAGTGACAGCAATGC

Alb-Cre_R: GACGGAAATCCATCGCTCGACCAGTTTAGT

Conditional lox excision was analyzed using following primers:

Excision_F: TCTGCTAACCATGTTCATGCCTTC

Excision_R: ATCGCACAGCTAGCGTTCGAG

### MITO-Tag Mice accessibility

MITO-Tag Mice (B6.Cg-*Gt(ROSA)26Sor^tm1(CAG-EGFP)Dmsa^*/J, The Jackson Laboratory, stock number: 032290) have been deposited to The Jackson Laboratory for use by the research community.

### Mouse studies

C57BL/6J mice were obtained from The Jackson Laboratory. Animal studies and procedures were conducted according to a protocol approved by the Institutional Animal Care and Use Committee (IACUC) at The Rockefeller University. All mice were maintained on a standard light-dark cycle with food and water *ad libitum*. For the fasting and refeeding experiment, 18 male mice (10 HA-MITO mice, 8 Control-MITO mice) were randomly allocated into either the fasted or refed groups, with 5 HA-MITO mice and 4 Control-MITO mice per group. The fasting and refeeding experiments described in this study were performed as follows: mice were fasted overnight for 12 h and then either refed or fasted for the next 4 h. Following cervical dislocation, liver tissue was immediately harvested for further processing. Note that one of the HA-MITO mice from the refed group was excluded from all analyses as this mouse did not exhibit the expected elevation in liver P-S6 signal upon refeeding, perhaps due to low food intake during the refeeding period (see Fig. 3*B* for expected P-S6 dynamics during fasted and refed conditions). The same mice were used for the S6 signaling immunoblot in Fig. 3*B*, the organellar marker immunoblot in Fig. S1, the lipidomic analysis, and the polar targeted and untargeted analyses. To avoid decreasing the speed of the workflow, immunopurifications intended for different downstream analyses were conducted in parallel for a given animal.

### Protein extraction from liver tissue

Liver tissue was excised using a punch biopsy tool and was washed three times in ice-cold PBS before treatment with detergent lysis buffer (10 mM Tris-Cl pH 7.5, 150 NaCl, 1 mM EDTA, 1% Triton X-100, 2% SDS, CHAPS 0.1%) supplemented with protease and phosphatase inhibitors. Homogenization was performed in the presence of cooling with liquid nitrogen with Bead Ruptor and the protein concentration was determined by using the Pierce BCA Protein Assay kit with bovine serum albumin as a protein standard.

### Immunofluorescence

Following cervical dislocation and dissection, 4 mm x 4 mm chunks of liver were cut and washed in ice-cold PBS. Next, the liver tissues were fixed in 10% neutral buffered formalin for 8 h, followed by washing three times in 70% ethanol. The histological preparations of the specimens in paraffin blocks were done in the Center of Comparative Medicine and Pathology, Memorial Sloan Kettering Cancer Center, New York. Liver sections were deparaffinized, rehydrated, and blocked (5% normal donkey serum, 0.3% Triton X-100 in PBS) for 2 h at room temperature. The sections were then incubated with primary antibodies (anti-HA, Cell Signaling Technology #3724 (1:100); anti-COX4l1, Abcam #ab14744 (1:100)) for 48 h at 4C. After washing, the sections were incubated with Alexa Fluor-conjugated secondary antibodies (Alexa Fluor 488-conjugated donkey anti-rabbit IgG (1:500); Alexa Fluor 568-conjugated donkey anti-mouse IgG (1:500)) for 2 h at room temperature, followed by counterstaining with Hoechst 33342 to label nuclei. Images were acquired on a RPI spinning disk confocal microscope using MetaMorph acquisition software.

### Rapid isolation of hepatocyte mitochondria from liver tissue

Isolation of mitochondria was performed as previously described with modifications (2, 4). All procedures were performed on ice. The liver tissue was excised with the punch biopsy tool on the cutting mat and was immediately washed three times in ice-cold PBS prior to extraction. Two pieces of tissue were homogenized in 5 ml of KPBS (136 mM KCl, 10 mM KH_2_PO_4_, pH 7.25 in Optima LC/MS water) with pestle attached to the mixer to rotate at 220 rpm. 5 μl of homogenate was taken as a sample of liver tissue. Homogenates were then spun down at 1000 x *g* for 2 min at 4°C. The supernatant was subjected to immunoprecipitation with pre-washed anti-HA beads for 3.5 min, followed by three rounds of washing in KPBS. In the last wash, 20% of the suspension of beads was set aside and used for protein extraction via detergent lysis. The remainder of the beads, as well as the liver tissue sample, were then extracted using a reagent appropriate for the desired downstream analysis.

### Immunoblotting

All liver tissue and isolated mitochondria samples were resolved on 8% or 12% SDS-PAGE gels and analyzed by immunoblotting as described previously (24).

### Proteomics

Proteins from lysates of isolated mitochondria were purified by trichloroacetic acid precipitation and the resulting pellet was resuspended in a 100 μl buffer of 6 M urea and 100 mM Tris at pH 7.8. Reduction and alkylation of disulfide bonds was then carried out by the addition of 5 μl 200 mM dithiothreitol (DTT) for 60 min to reduce disulfide bonds. The resulting free cysteine residues were subjected to an alkylation reaction by the addition of 20 μl 200 mM iodoacetamide for 60 min to form carbamidomethyl cysteine. This solution was brought to a volume of 900 μl to reduce the urea concentration. 100 μl of 20 ng/μl of trypsin was added for a final volume of 1000 μl. These were allowed to digest overnight at 37°C with gentle shaking. The resulting peptides were washed, extracted, and concentrated by solid phase extraction using Waters Sep-Pak Plus C18 cartridges. Organic solvent was removed and the volumes were reduced to 20 μl using a speed vac for subsequent analyses.

For chromatographic separation, the digestion extracts were analyzed by reversed phase high performance liquid chromatography (HPLC) using Thermo EASY-nLC 1200 pumps and autosampler and a Thermo Q Exactive HF-X Hybrid Quadrupole-Orbitrap mass spectrometer using a nano-flow configuration. Samples were loaded on a 6 cm x 100 μm column packed with 10 μm ODS-A C18 material (YMC) and washed with 4 μl total volume to trap and wash peptides. These were then eluted onto the analytical column, which was self-packed with 1.7 μm Aeris C18 material (Phenomenex) in a fritted 14 cm x 75 μm fused silica tubing pulled to a 5 μm tip. The gradient was isocratic 1% Buffer A for 1 min at a rate of 250 nL/min with increasing buffer B concentrations to 19% buffer B at 52 min, 34% buffer B at 77.5 min, and 50% buffer B at 103 min. The column was washed with high percent buffer B and re-equilibrated between analytical runs for a total cycle time of approximately 127 min. Buffer A consisted of 1% formic acid in water and buffer B consisted of 1% formic acid in acetonitrile. The mass spectrometer was operated in a dependent data acquisition mode where the 20 most abundant peptides detected in the Orbitrap using full scan mode with a resolution of 60,000 were subjected to daughter ion fragmentation in the linear ion trap. A running list of parent ions was tabulated to an exclusion list to increase the number of peptides analyzed throughout the chromatographic run.

Peptides were identified from the mass spectrometry (MS) data using PEAKS Studio 8.5. The *mus musculus* Refseq protein FASTA entries were downloaded from NIH/NCBI and concatenated to a database of common contaminants (keratin, trypsin, etc). An FDR threshold of 1% for identification of peptides was used.

### Lipidomics

To extract lipids, 333 μL of 100% methanol was added to the anti-HA beads or liver tissue sample, followed by 667 μL of chloroform. Samples were vortexed at max speed in a cold room for 15 min, followed by the addition of 200 μl of 0.9% (w/v) NaCl. Lipids were vortexed at 4°C for 15 min and were spun down at 3000 x *g* for 15 min at 4°C. The bottom layer containing the lipids were taken and briefly dried under a cold stream of nitrogen gas.

Lipids were separated on an Ascentis Express C18 2.1 × 150mm 2.7 um column (MilliporeSigma) connected to a Dionex UlitMate 3000 UPLC system and a QExactive benchtop orbitrap mass spectrometer (Thermo Fisher Scientific) equipped with a heated electrospray ionization (HESI) probe. Dried lipid extracts were reconstituted in 50 μL 65:30:5 acetonitrile: isopropanol: water (v/v/v). 5 μL of sample were injected onto the column, with separate injections for positive and negative ionization modes. Mobile phase A in the chromatographic method consisted of 60:40 water: acetonitrile with 10 mM ammonium formate and 0.1% formic acid, and mobile phase B consisted of 90:10 isopropanol: acetonitrile, with 10 mM ammonium formate and 0.1% formic acid. The chromatographic gradient was described previously (25). The column oven and autosampler were held at 55°C and 4°C, respectively. The mass spectrometer parameters were described previously (26) and were modified as described in (27). The spray voltage was set to 4.2 kV, and the heated capillary and the HESI were held at 320°C and 300°C, respectively. The S-lens RF level was set to 50, and the sheath and auxillary gas were set to 35 and 3 units, respectively. These conditions were held constant for both positive and negative ionization mode acquisitions. External mass calibration was performed every 7 days using the standard calibration mixture.

Mass spectra were acquired in both full-scan and data-dependent MS/MS mode. For the full-scan acquisition, the resolution was set to 70,000, the AGC target was 1×10^6^, the maximum injection time was 50 ms, and the scan range was *m/z* = 133.4 - 2000. For data-dependent MS/MS, the top 10 ions in each full scan were isolated with a 1.0-Da window, fragmented with a step-wise collision energy of 15, 25, and 35 units, and analyzed at a resolution of 17,500 with an AGC target of 2×10^5^ and a maximum injection time of 100 ms. The underfill ratio was set to 0. The selection of the top 10 ions was set to isotopic exclusion, a dynamic exclusion window of 5.0 s, and an exclusion list of background ions based on a solvent bank.

High-throughput identification and relative quantification of lipids was performed separately for positive and negative ionization mode data, using LipidSearch software (ThermoFisher Scientific/ Mitsui Knowledge Industries) (28, 29) using the default parameters for QExactive product search and alignment. After alignment, raw peak areas for all identified lipids were exported to Microsoft Excel and filtered according to the following predetermined quality control criteria: Rej (“Reject” parameter calculated by LipidSearch) equal to 0; PQ (“Peak Quality” parameter calculated by LipidSearch software) greater than 0.85; CV (standard deviation/ mean peak area across triplicate injections of a represented (pooled) biological sample) below 0.4; *R* (linear correlation across a three-point dilution series of the representative (pooled) biological sample) greater than 0.9. Typically, ~70% of identified lipids passed all four quality control criteria. Cardiolipins were validated by manually checking the peak alignment and matching the MS/MS spectra to the characteristic fragmentation patterns found in the LIPIDMAPS^®^ database (www.lipidmaps.org), as well as those described previously (30). Validated cardiolipin species were reanalyzed using Xcalibur Quanbrowser 4.1 (Thermo Fisher Scientific), and graded based on the quality of MS/MS spectra collected: Grade A - CL annotation is correct, all four acyl chains are correctly annotated; Grade B - 3/4 acyl chains are correctly annotated; Grade C - 2/4 acyl chains are correctly annotated; Grade D - 1/4 acyl chains are correctly annotated or CL species confirmed through presence of head group, but acyl chain identification is ambiguous.

For a lipid species to be examined for mitochondrial abundance, it had to pass the aforementioned four quality control criteria. For cardiolipin analysis, only grade A cardiolipins were used. A lipid species was only considered mitochondrial within a given dietary state if 1) the mean Control-MITO IP peak area was 0 and the peak areas of each HA-MITO IP replicate were greater than 0, or 2) the mean Control-MITO IP peak area was greater than 0 but the peak areas of each HA-MITO IP replicate were at least 1.5-fold the mean Control-MITO IP peak area.

### Polar metabolomics

For polar metabolites, the extraction of liver tissue or isolated mitochondria was performed using 80% ice-cold methanol with internal standards. Metabolite profiling was conducted on a QExactive bench top orbitrap mass spectrometer equipped with an Ion Max source and a HESI II probe, which was coupled to a Dionex UltiMate 3000 HPLC system (Thermo Fisher Scientific, San Jose, CA). External mass calibration was performed using the standard calibration mixture every 7 days. Typically, 5 μL were injected onto a SeQuant^®^ ZIC^®^-pHILIC 150 × 2.1 mm analytical column equipped with a 2.1 × 20 mm guard column (both 5 mm particle size; EMD Millipore). Buffer A was 20 mM ammonium carbonate, 0.1% ammonium hydroxide; Buffer B was acetonitrile. The column oven and autosampler tray were held at 25°C and 4°C, respectively. The chromatographic gradient was run at a flow rate of 0.150 mL/min as follows: 0-20 min: linear gradient from 80-20% Buffer B; 20-20.5 min: linear gradient form 20-80% Buffer B; 20.5-28 min: hold at 80% Buffer B. The mass spectrometer was operated in full-scan, polarity-switching mode, with the spray voltage set to 3.0 kV, the heated capillary held at 275°C, and the HESI probe held at 350°C. The sheath gas flow was set to 40 units, the auxiliary gas flow was set to 15 units, and the sweep gas flow was set to 1 unit. MS data acquisition was performed in a range of *m/z* = 70-1000, with the resolution set at 70,000, the AGC target at 1×10^6^, and the maximum injection time at 20 ms. A sample of KPBS in 80% methanol with internal standards was included in each polar metabolomics run to monitor for signs of buffer contamination. Relative quantitation of polar metabolites was performed with XCalibur QuanBrowser 4.1 (Thermo Fisher Scientific) using a 5 ppm mass tolerance and referencing an in-house library of chemical standards. Peak area ratios were calculated using the peak area of each metabolite and the peak area of an internal standard. No adjustment for KPBS matrix effects was done for polar metabolomics as we only performed relative quantification and all samples had KPBS in them.

Determination of mitochondrial abundances of polar metabolites within hepatocytes during fasted and refed conditions was done as follows. Within a given dietary state, metabolites were classified as mitochondrial if 1) the mean Control-MITO IP peak area ratio was 0 and the peak area ratios of each HA-MITO IP replicate were greater than 0, or 2) the mean Control-MITO IP peak area ratio was greater than 0 but the peak area ratios of each HA-MITO IP replicate were at least 1.5-fold the mean Control-MITO IP peak area ratio. Only metabolites that were deemed mitochondrial were then used for further calculations. Within a given dietary state, background correction was done for each mitochondrial metabolite by subtracting the mean Control-MITO IP peak area ratio from the peak area ratio of each HA-MITO IP replicate. Background-corrected HA-MITO IP metabolite values were then normalized based on the amount of isolated mitochondria in each HA-MITO IP, which can be assessed via immunoblotting for citrate synthase and quantitation of band intensities. (Note that in this work, we found that the background-corrected HA-MITO IP peak area ratio of the mitochondrial metabolite FAD actually performed better than citrate synthase for estimating the amount of isolated mitochondria in each HA-MITO IP and was thus used for normalizing samples.) Because of the reproducibility of the punch biopsy tool, equal amounts of liver tissue were homogenized and then taken for metabolite extraction and so no normalization was done for the liver tissue samples.

### Untargeted polar metabolomics

Data were acquired as described above, with additional data-dependent (dd) MS/MS collected on pooled samples to aid with unknown metabolite identification. For ddMS/MS, the top 10 ions in each full scan were isolated with a 1.0-Da window, fragmented with a step-wise collision energy of 15, 30, and 45 units, and analyzed at a resolution of 17,500 with an AGC target of 2×10^5^ and a maximum injection time of 100 ms. The underfill ratio was set to 0. The selection of the top 10 ions was set to isotopic exclusion, a dynamic exclusion window of 5.0 s, and an exclusion list of background ions based on a solvent bank. Data were analyzed using Compound Discoverer 2.1 (Thermo Fisher Scientific) and by including an in-house mass-list. P values were adjusted according to the Benjamini-Hochberg method. O-adipoylcarnitine, O-succinylcarnitine, 4-guanidinobutyric acid, methionine, and alanine were validated using chemical standards. These metabolites were also analyzed in a targeted fashion. All analyses and calculations for these metabolites were done as described previously for polar mitochondrial metabolites within hepatocytes (see “Polar Metabolomics” section).

### Statistics

All p values were calculated using an unpaired, two-tailed, parametric student’s *t* test using Graph Pad Prism 7 except, for Fig. 4*A* and Table S4, p values were calculated using Compound Discoverer 2.1 and then adjusted using the Benjamini-Hochberg method.

## AUTHOR CONTRIBUTIONS

E.C.B., D.M.S., K.B., and W.W.C. initiated the project and designed the research. E.C.B. conducted most of the experimental work and W.W.C. performed most of the data analysis. L.B. and C.Ö. assisted E.C.B. with experimentation. C.A.L., S.H.C., and T.K. operated the LC/MS equipment. C.A.L. and S.H.C. provided the raw data for all metabolomics work. C.A.L., S.H.C., and W.W.C. analyzed the metabolomics data. M.A.R. assisted E.C.B. with carrying out the proteomic work and assisted W.W.C. with analyzing the proteomic data. A.L.C. assisted E.C.B. with performing the immunofluorescence. E.C.B., D.M.S., K.B., and W.W.C. wrote and edited the manuscript.

## ACKNOWLEDGMENTS

We thank all members of the Birsoy and Sabatini laboratories and Elizaveta Freinkman and Hoi See Tsao for helpful suggestions and thank members of the Boston Combined Residency Program for their assistance. C4- and C5-carnitines were kindly synthesized by Rajan Pragani and provided by Jared Mayers. E.C.B is supported by Robertson Therapeutic Funds of The Rockefeller University. A.L.C. is supported by an NIH F31 NRSA fellowship. K.B. is supported by DP2 (DP2 OD024174-01), Irma-Hirschl Trust, AACR NextGen Grant, and is a Searle Scholar, Sidney Kimmel Scholar, Pew-Stewart Scholar and Basil O’Connor Scholar of March of Dimes. This work was supported by grants from the US NIH (R01CA103866, R01CA129105, and R37AI047389) and the Department of Defense (W81XWH-15-1-0230) to D.M.S., and D.M.S. is an investigator of the Howard Hughes Medical Institute and an American Cancer Society Research Professor. W.W.C. is supported by the Boston Combined Residency Program and Boston Children’s Hospital.

---

**Fig. S1.**
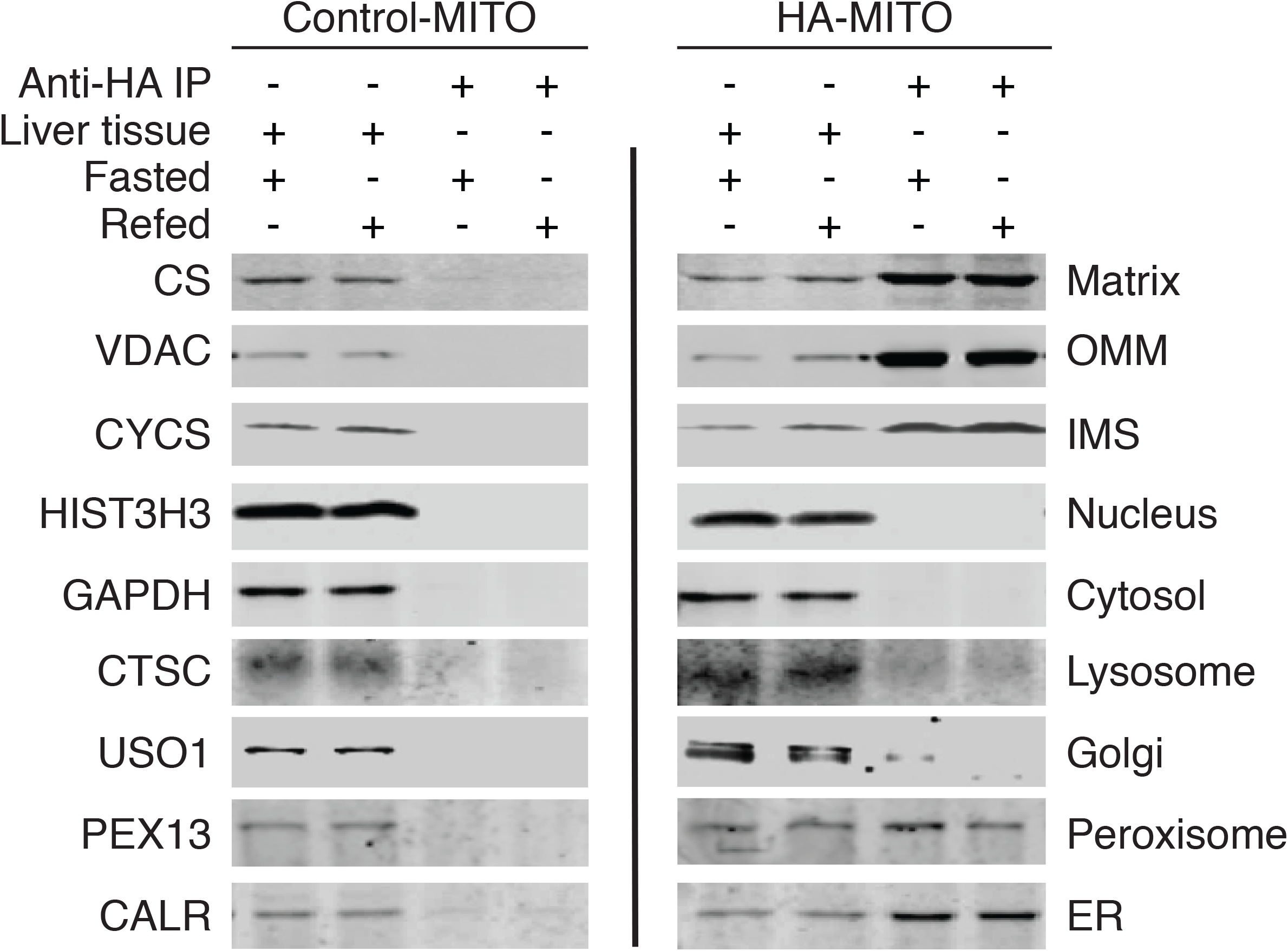
Characterization of IP purity for mitochondrial isolations performed on mice during fasted and refed conditions. Immunoblot analysis of liver tissue and the anti-HA IPs from Control-MITO and HA-MITO mice. The names of the protein markers used and the corresponding subcellular compartments appear to the left and right of the immunoblots, respectively. Matrix, mitochondrial matrix; OMM, outer mitochondrial membrane; IMS, mitochondrial intermembrane space; Golgi, Golgi complex; ER, endoplasmic reticulum; Control-MITO, Control-MITO mice; HA-MITO, HA-MITO mice.

